# SLAM-ITseq: Sequencing cell type-specific transcriptomes without cell sorting

**DOI:** 10.1101/235093

**Authors:** Wayo Matsushima, Veronika A Herzog, Tobias Neumann, Katharina Gapp, Johannes Zuber, Stefan L Ameres, Eric A Miska

## Abstract

Cell type-specific transcriptome analysis is an essential tool in understanding biological processes but can be challenging due to the limits of microdissection or fluorescence-activated cell sorting (FACS). Here, we report a novel *in vivo* sequencing method, which captures the transcriptome of a specific type of cells in a tissue without prior cellular or molecular sorting. SLAM-ITseq provides an accurate snapshot of the transcriptional state *in vivo*.

---

Animals consist of various organs, which are further composed of heterogeneous populations of highly specialised cells. Thus, it is important to look at transcriptomic changes at the cellular level to understand animal physiology. To capture the transcriptome of a specific cell type, mechanical cell isolation methods such as fluorescence-activated cell sorting (FACS) or laser capture microdissection (LCM) prior to RNA extraction have widely been used so far. Moreover, combined with such cell isolation methods, the recent advance in high-throughput RNA sequencing (RNA-seq) methods now enables us to quantitate transcripts at single-cell resolution ^1^. However, since the transcriptome of a cell is greatly affected by its cellular context as well as mechanical/chemical stimuli, it has been questioned how closely transcriptomic data obtained from sorted cells reflect the status of before cell sorting ^2^. Also, these cell isolation methods are often time-intensive, involve laborious steps and lead to considerable cell death after isolation, which limits their applications to robust cells only.

Recently, Gay *et al.* have developed an elegant *in vivo* metabolic RNA labeling method, to study cell type-specific transcriptomes, using a uracil analogue 4-thiouracil ^3,4^. *Toxoplasma gondii* (*T. gondii*) uracil phosphoribosyltransferase (UPRT) converts 4-thiouracil to uridine monophosphate (UMP) and then incorporates it into newly synthesised RNA molecules to generate thiol-containing RNA (thio-RNA), while UPRT homologs in many organisms, including mice, are enzymatically inactive. Gay *et al.* generated transgenic mice expressing *T. gondii* UPRT in specific cell type and exposed them to 4-thiouracil to“label” newly synthesised RNA. Then, the thio-RNA was pulled-down by a biochemical isolation method and quantified by RNA-seq to identify its enrichment over the total RNA. Even though similar methods have been tested in various model organisms ^5–7^, technical and analytical challenges limit its application. First, the biochemical isolation methods of thio-RNA have been shown to have high background noise, which makes it difficult to identify lowly labelled transcripts and to distinguish the labelled RNA from background noise. This issue is especially pronounced when used *in vivo,* where relatively low 4-thiouracil concentrations can be achieved. Second, because the labelled and unlabelled population of RNA are sequenced separately, unbiased estimation of the labelling level of a given transcript is difficult. Also, to determine whether a given transcript is labelled, the enrichment level of the pulled-down read count over the input read count was used in this original method. However, this estimation is not optimal for tissues where UPRT-expressing cells are the dominant cell type, as the read count between the pulled-down and input fractions are mostly the same, and thus it is hard to determine which transcripts are labelled.

Here, we describe a new method that significantly improves *in vivo* metabolic labelling: we redesigned the experiment to use a different control to account for background labelling, used the RNA-seq method called thiol(SH)-linked alkylation for the metabolic sequencing of RNA (SLAMseq) to directly identify thiol-containing uracil at single-base resolution ^8^, and applied a statistical method to reliably identify labelled transcripts, accounting for biological variance in the labelling level. This improved method, which we now term SLAMseq *i*n *t*issue (SLAM-ITseq), makes the *in vivo* 4-thiouracil-based metabolic labelling methods accessible to wider research areas to study cell type-specific transcriptomics in animals.

First, to compare SLAM-ITseq to the previously described isolation method, we generated the same double-transgenic mice used, which label RNA in endothelial cells specifically. Homozygous *UPRT* transgenic mice (*uprt/uprt*), which have floxed stop codons and SV40 polyadenylation sequences upstream of the UPRT-coding transgene, were crossed with hemizygous *Tie2-Cre* mice (*cre/0*) ^9^, and then *uprt/0; cre/0* (Cre+) and *uprt/0*; +/+ (Cre-) animals were obtained. Since the Tie2 promoter is known to be specific to endothelial cells in the brain, UPRT is expressed only in this specific cell type in Cre+ animals (Supplementary Fig. 1a).

To control for the background labelling and to capture both specific and common transcripts of a certain cell type, we used the Cre-animals as a control. After exposing both Cre+ and Cre-animals to 4-thiouracil for 4 hours, RNA was extracted from the whole brain from each animal (Fig. 1a). To confirm controlled *UPRT* transgene expression, reverse transcription followed by quantitative polymerase chain reaction (RT-qPCR) was performed on cDNA obtained from each animal. As expected, it was confirmed that UPRT was expressed only in Cre+ animals (Fig. 1b). To identify the labelled transcripts, the remaining RNA was treated with iodoacetamide (IAA) to alkylate the thiol group of the thio-RNA and then subsequently used as RNA-seq input. During the reverse transcription step of RNA-seq library preparation, a guanine (G), instead of adenine (A), is base-paired to an alkylated 4-thiouracil leading to the thymine to cytosine base conversion (T>C) at the corresponding T position in the reads generated from the thio-RNA. T>C mismatch-aware alignment and T>C count per gene were performed using the software SLAM-DUNK. As expected, significantly more T>Cs were observed in Cre+ animals (Fig. 1c). Importantly, the transcriptome itself remained unchanged between Cre+ and Cre-animals (Supplementary Fig. 1b), suggesting that both Cre and UPRT expression has little effects on the overall transcriptome.

**Figure 1.**
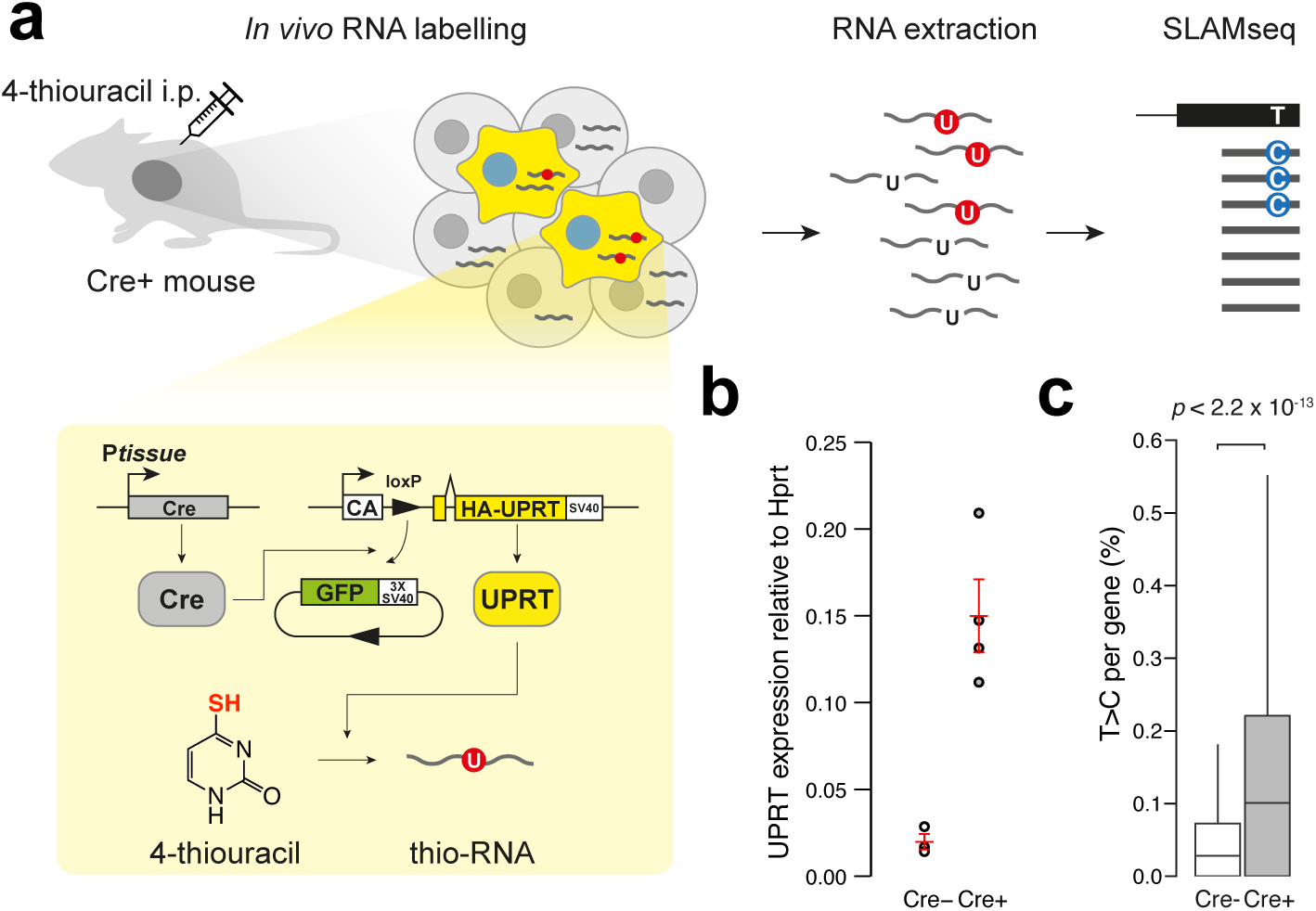
SLAM-ITseq design. (a) Schematic of how SLAM-ITseq works. P*tissue*, a tissue-specific promoter; CA, chicken beta-actin promoter. (b) Comparison of UPRT mRNA expression by RT-qPCR in total brain RNA from Cre+ and Cre-animals. The red bars indicate the mean expression and 95% confidence intervals (Cre+: n=4, Cre-: n=3). (c) Comparison of T>C rate in all T positions sequenced per gene. Data are shown as boxplots. The lower and upper hinges correspond to the first and third quartiles, the middle hinges indicate the median, and the whiskers extend to 1.5 interquartile range from the upper hinges. Outliers are not shown. Mann-Whitney *U* test was used to calculate the *p*-value presented.

Next, we employed a statistical approach to identify the labelled transcripts. Since nucleotide conversion is a binomial process and the probability of it occurring at a given T position can be modelled by beta distribution for biological replicates, beta-binomial distribution can describe the T>C fraction per gene among biological replicates. Based on the variance estimate of T>C fraction, genes that are significantly labelled in Cre+ were identified at FDR < 0. 05 (Fig. 2a). To evaluate the sensitivity of this significance calling, we used the same sets of known endothelial genes selected by Gay *et al.* as a positive control. As expected, out of 13 known endothelial genes (*Cdh5, Cd34, Egfl7, Emcn, Esam, Ets1, Flt1, Kdr, Nos3, Pecaml, Tek, Tie1, and Thsdl*), 11 were called as significant (Fig. 2b), except for *Tie1* and *Cd34,* which were not called as significant in the Gay *et al.* paper as well. Also, among a few known neuronal genes examined, no significantly labelled ones were observed (Fig. 2c). Together, these results indicate successful labelling of endothelial transcripts without labelling the transcripts in surrounding neuronal cells. To comprehensively analyse the functional profile of significantly labelled genes, gene ontology (GO) terms enrichment analysis was performed on the labelled gene list. The biological process GO terms in the list were revealed to enrich for GO terms that are well known for endothelial functions, such as “vasculature development” or “angiogenesis” (Fig. 2d). These results confirm a statistical enrichment for transcripts synthesised in endothelial cells. Interestingly, the labelled gene list also contained known housekeeping genes such as *Hprt* and *Actb* (Supplementary Fig. 1c), which should be expressed in both the endothelial cells and the other cell types, despite the majority being unlabelled. Collectively, these results show that this improved method enables unbiased capture of the transcriptome of a cell type of interest, including commonly expressed genes, which was not possible with the original method.

**Figure 2.**
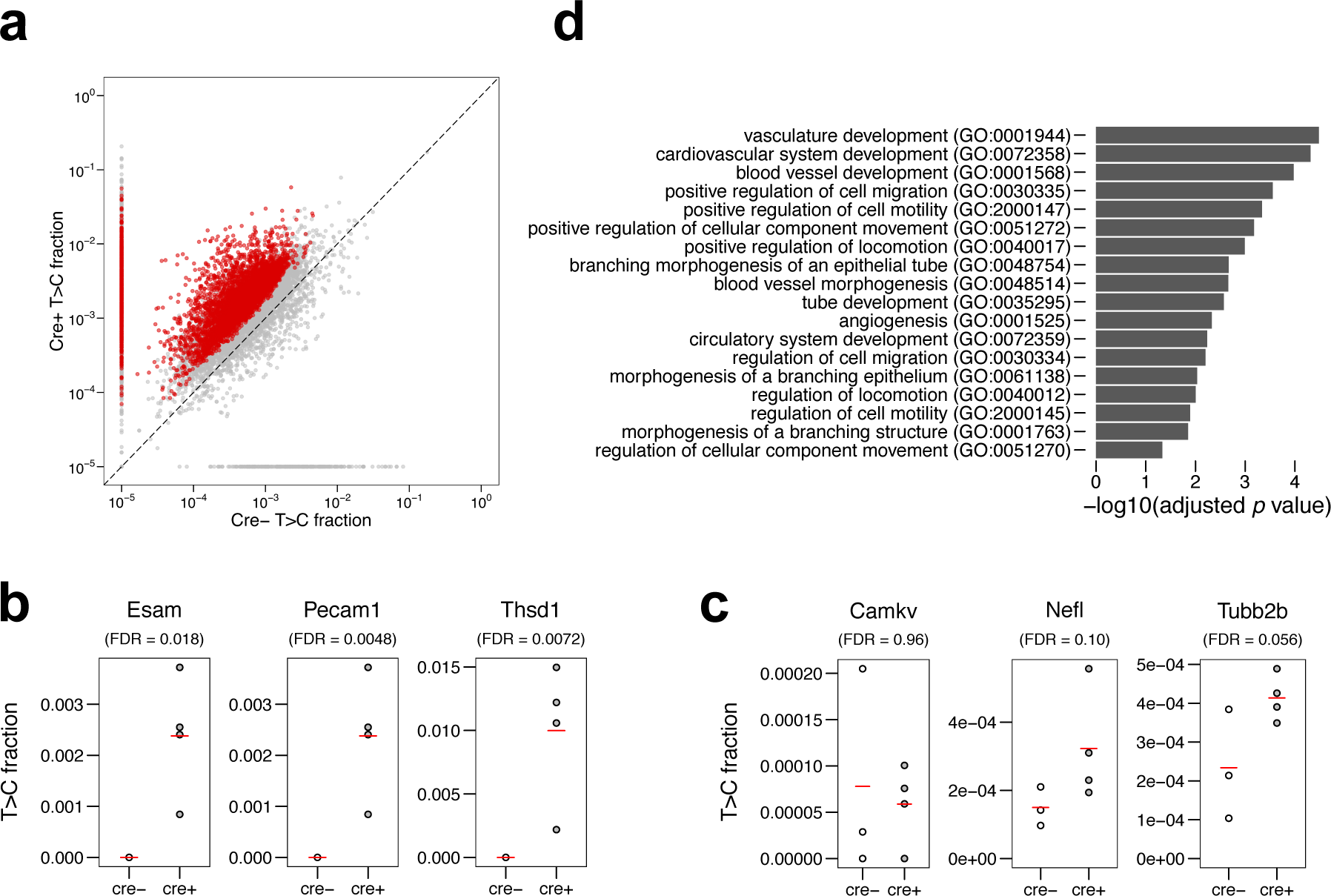
Analyses of labelled RNA from the mouse brain expressing UPRT in endothelial cells. (a) Average T>C fractions of each gene are plotted. Significantly more labelled transcripts in Cre+ were determined by beta-binomial test and shown as red points (FDR < 0.05). A constant value of 1x10^-5^ was added to the row T>C value when plotting. (b, c) T>C fraction of known endothelial cell-specific genes (b) and neuronal cell-specific genes (c) are shown. The red bars indicate the mean T>C fraction of biological replicates (Cre+: n=4, Cre-: n=3). (d) GO term enrichment analysis performed on top 130 significantly labelled transcripts.

Next, we examined if this method is applicable to study transcriptomes of other cell types, using different promoters for the Cre expression control. *Vil-Cre* ^10^ and *Adipoq-Cre* ^11^ mice were crossed with the *UPRT* mice to generate mice that specifically express UPRT in adipocytes and gut epithelial cells, respectively. It is important to note that the ratio of Vil+ cells in intestine and Adipoq+ cells in adipose tissue are much higher than Tie2+ cells in the brain, and thus it was difficult to make use of such Cre lines with the conventional method. RNA was extracted from epididymal white adipose tissue (eWAT) and duodenum from each set of mice. Cre-dependent UPRT expression was confirmed by RT-qPCR (Supplementary Fig. 2b, 3b). To identify the labelled RNA, SLAMseq was performed on the RNA extracted from each tissue. Beta-bionomial test revealed that genes known to be expressed in the Cre-expressing cells were significantly labelled, while those known to be expressed in the other cell types in the tissue were not labelled (Supplementary Fig. 2e-h, 3e,f). These results demonstrate that SLAM-ITseq can identify cell type-specific transcripts using Cre lines specific to a wide range of tissues, regardless of the proportion of the cell type of interest in a tissue.

SLAM-ITseq dramatically broadens the application of SLAMseq to the study of cell type-specific transcriptomes *in vivo* from a wide range of tissues without cell isolation. Thus, SLAM-ITseq provides unparalleled access to cellular transcriptional dynamics to better understand animal physiology.

## Methods

### Animal husbandry

All mice were maintained in a specific pathogen free facility with sentinel monitoring at standard temperature (19-23°C) and humidity (55% ±10%), on a 12h dark, 12h light cycle (lights on 0730–1900) and fed a standard rodent chow (LabDiet 5021-3, 9% crude fat content, 21% kcal as fat, 0.276ppm cholesterol, LabDiet, London, UK). Both food and water were available *ad libitum*. The mice were housed in groups of 3-4 mice per cage in individually ventilated caging receiving 60 air changes per hour. In addition to Aspen bedding substrate, standard environmental enrichment of a nestlet and a cardboard tunnel were provided. All animals were regularly monitored for health and welfare concerns and were additionally checked prior to and after procedures. The care and use of mice in the study was carried out in accordance with UK Home Office regulations, UK Animals (Scientific Procedures) Act of 1986 under a UK Home Office license that approved this work (PF8733E07), which was reviewed regularly by the WTSI Animal Welfare and Ethical Review Body.

### Generation of transgenic mice

Homozygous *UPRT* transgenic mice ^4^ (*uprt/uprt*) were crossed with hemizygous *Cre* mice (*Tie2-Cre* : JAX stock #008863, *Vil-Cre* : JAX stock #004586, and *Adipoq-Cre* : JAX stock #010803) ^9–11^ *(cre/0),* and then *uprt/0; cre/0* (Cre+) and *uprt/0*; +/+ (Cre-) animals were obtained as resulting offspring. To confirm the genotype, DNA from mouse ear-clips was isolated using the Sample-to-SNP kit (Life Technologies), and products amplified using a Viia7 qPCR machine (Life Technologies). Results were analysed against known calibrator controls via the ddCt method ^12^. TaqMan (Life Technologies) qPCR assay sequences against UPRT are: Forward 5’-ATTCCAAGATCTGTGGCGTC-3’, Reverse 5’-CTTCTCGTAGATCAGCTTAGGC-3’, Probe (VIC) 5’-CCGCATCGGGAAAATCCTCATCCA-3’, and against Cre are: Forward 5’-ACGTACTGACGGTGGGAGAA-3’, Reverse 5’-GTGCTAACCAGCGTTTTCGTT-3’, Probe (VIC) 5’-CTGCCAATATGGATTAACA-3’.

### 4-thiouracil administration

4-thiouracil administration was performed following the previously reported methods ^4,13^. Briefly, 4-thiouracil was dissolved in DMSO at 200 mg/ml concentration followed by further dilution in corn oil in 1:4 ratio. This solution was intraperitoneally injected to 8-10 week old Cre+ and Cre-mice using a 25G x ⅝” needle. 4 hours later, mice were culled and tissues were harvested. The collected tissues were cut into small pieces (less than 5 mm in thickness) and submerged in RNAlater (Sigma-Aldrich) for storage at −20C.

### RNA extraction from tissues

After removing RNAlater by clean Kimtech (Kimberly-Clark), around 30 mg of tissue was homogenised in 1 ml TRIsure (Bioline), using TissueLyser LT (Qiagen) and 7 mm Stainless Steel Beads (Qiagen). RNA extraction was performed following the manufacturer’s instruction with minor modifications: the isopropanol precipitation step was done with the addition of 20 ug/ml glycogen and 0.1 mM DTT and the isopropanol precipitation at −20C was extended to 2 hr. Purified RNA was subsequently treated with 1 U of TURBO DNase (Invitrogen) to eliminate potential DNA contamination. After the DNase reaction, RNA was cleaned up by RNA Clean & Concentrator™-5 (Zymo Research) and stored at −80C with 1 mM DTT in order to prevent oxidation.

### RT-qPCR

Reverse transcription was performed by SuperScript II Reverse Transcriptase (Invitrogen) and random hexamers (Invitrogen) following the manufacturer’s instruction. Around 10 μg of the synthesised cDNA was used as an input for 10 μl qPCR reaction using PowerUp SYBR Green Master Mix (Applied Biosystems) with the gene specific primer pairs targeting *T. gondii* UPRT (Forward: 5’-CCCGATATTCGACAAACGAC-3’, Reverse: 5’-GCTTCATGAGCACCACATTG-3’) and *M. musculus* Hprt (Forward: 5’-GCCTAAGATGAGCGCAAGTTG-3’, Reverse: 5’-TACTAGGCAGATGGCCACAGG-3’). Technical triplicate was prepared and the mean CT value was calculated for each biological replicate.

### SLAMseq

DNase-treated RNA was reacted with IAA to alkylate the thiol group following the protocol previously described ^14^ Briefly, 50 μl reaction mix (5-10 μg RNA, 10 mM IAA, 50 mM pH8 sodium phosphate, and 50% DMSO) was incubated at 50C for 15 min. The reaction was stopped by adding 1 μl of 1M DTT, followed by 1 μl glycogen (20 mg/ml), 5 μl NaOAc (3M, pH 5.2), and 125 μl 100% EtOH addition. After 2 hr incubation at −20C, the solution was centrifuged and the obtained RNA pellet was washed with 80% ethanol. The pellet was resuspended in 15 ul nuclease-free water. RNA concentration was quantified by Qubit RNA BR Assay Kit (Molecular probes) and 500 ng of it was used as an input for QuantSeq 3' mRNA-Seq Library Prep Kit for Illumina (Lexogen). RNA-seq library preparation was conducted following manufacturer’s instructions with the PCR cycles optimised for different tissues (brain: 13, duodenum: 15, eWAT: 15). The multiplexed libraries were sequenced using HiSeq 1500 (Illumina) for single-end 100 cycles by the core NGS service at the Gurdon Institute.

### Bioinformatics

Demultiplexed fastq files were first analysed with FastQC (version 0.11.5) for quality check. Then, these sequence data were analysed by the software designed for SLAMseq analysis, SLAM-DUNK (version 0.2.4, http://t-neumann.github.io/slamdunk/), to quantify how many T>Cs are detected per gene in each sample. For the mapping and SNP calling, mouse primary genome assembly GRCm38 was used. For counting T>C per gene, 3’UTR annotation data from Refseq and Ensembl was used. The other parameters used were as following: −5 12 -n 100 -m -mv 0.2 -mts -rl 100.

To identify significantly labelled transcripts, first, genes that had zero coverage on T in any sample were excluded. Next, the number of T>Cs and the total T coverage of the genomic sequence in each annotated gene were used to perform beta-binomial test by the R package ibb (version 13.06) ^15^. To control the false discovery rate (FDR), the Benjamini-Hochberg procedure was employed on the calculated *p*-value, and FDR < 0.05 was set to determine the significantly labelled transcripts.

For GO term enrichment analysis, PANTHER 12.0 (http://pantherdb.org/) was used to perform the “Statistical overrepresentation test” on the top 130 significantly labelled transcripts.

### Statistics

Statistic analyses described and plots generation were performed using R (version 3.3.2).

## Data availability statement

Sequencing data and metadata associated with this manuscript is available at ArrayExpress (https://www.ebi.ac.uk/arrayexpress/) under the accession number E-MTAB-6353.

## Acknowledgement

We thank Kay Harnish for high-throughput sequencing support, Brian Reichholf and Pooja Bhat for technical support, and Wellcome Trust Sanger Institute Research Support Facility Staffs for mouse maintenance and experimental supports.

This work was supported by Cancer Research UK (C13474/A18583, C6946/A14492) and the Wellcome Trust (104640/Z/14/Z, 092096/Z/10/Z) to E.M. and the European Research Council (ERC-StG-338252 miRLIFE) to S.L.A. The IMP is generously supported by Boehringer Ingelheim. W.M. was supported by The Nakajima Foundation and St John’s College Benefactors’ Scholarship. K.G. was supported by Swiss National Foundation postdoc mobility fellowship.

## Author contributions

W.M. and E.A.M. conceived and designed the study and wrote the manuscript; W.M. and V.A.H. performed the experiments; W.M and T.N. analysed the data; K.G. advised on the study design; S.L.A., J.Z. and E.A.M. provided expertise and feedback.

---

**Figure S1.**
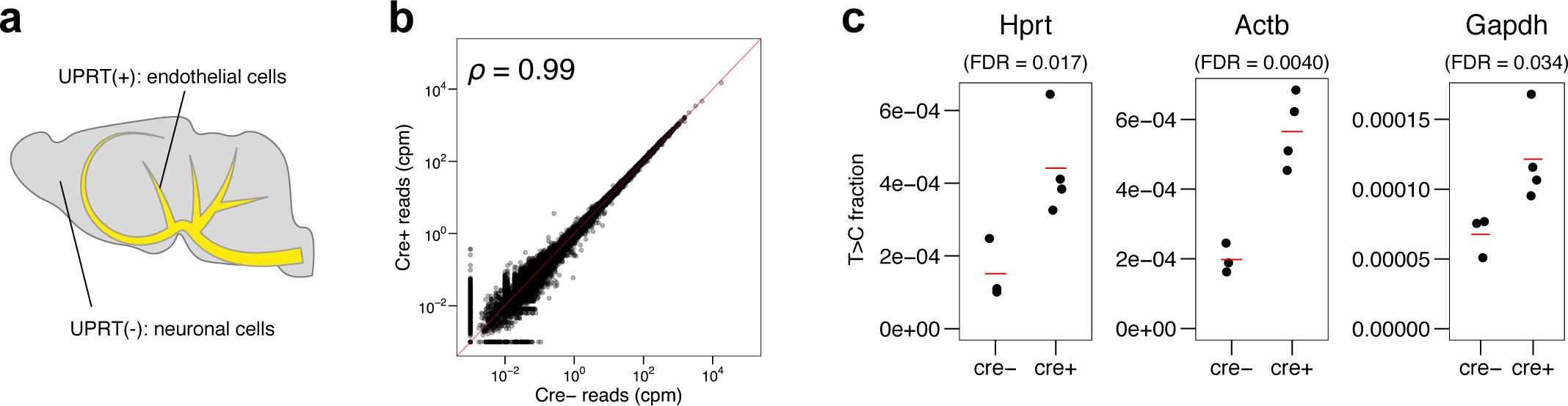
Additional analyses of SLAM-ITseq on endothelial cells in the brain. (a) Schematic of UPRT-expressing cells (yellow) and non-UPRT-expressing cells (grey) in the Cre+ mouse brain. (b) Comparison of RNA expression level between Cre+ and Cre-mice. Spearman’s correlation coefficient *ρ* is shown. (c) The labelling levels of the genes known to be expressed in both endothelial cells and other cells are shown. The red bars indicate the mean T>C fraction of biological replicates (Cre+: n=4, Cre-: n=3).

**Figure S2.**
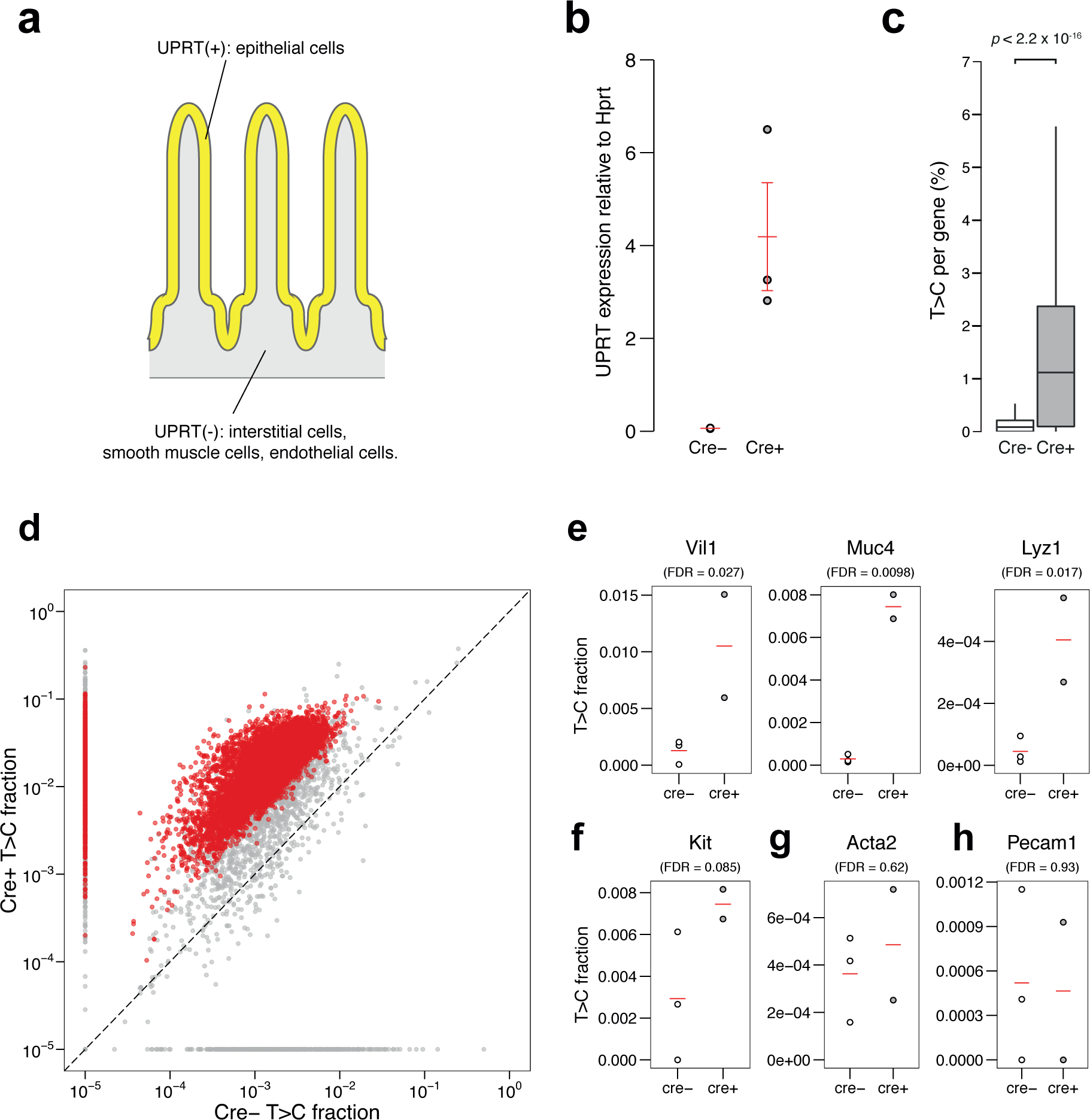
SLAM-ITseq analyses of labelled RNA from the mouse duodenum expressing UPRT in epithelial cells. (a) Schematic of UPRT-expressing cells (yellow) and non-UPRT-expressing cells (grey) in the Cre+ mouse intestine. (b) Comparison of UPRT mRNA expression by RT-qPCR in total duodenum RNA from the Cre+ and Cre-animals. The red bars indicate the mean expression and 95% confidence intervals (Cre+: n=2, Cre-: n=3). (c) Comparison of T>C rate in all T positions sequenced per gene. Data are shown as boxplots. The lower and upper hinges correspond to the first and third quartiles, the middle hinges indicate the median, and the whiskers extend to 1.5 interquartile range from the upper hinges. Outliers are not shown. Mann-Whitney *U* test was used to calculate the p-value indicated. (d) Average T>C fractions of each gene are plotted. Significantly more labelled transcripts in Cre+ were determined by beta-binomial test and shown as red points (FDR < 0.05). A constant value of 1x10^-5^ was added to the row T>C value when plotting. (e) T>C fraction of known intestinal epithelium-specific genes. (f-h) Genes known to be transcribed in non-epithelial cells in small intestine; an interstitial gene (f), a smooth muscle gene (g), and an endothelial gene (h) are shown. The red bars indicate the mean T>C fraction of biological replicates (Cre+: n=2, Cre-: n=3).

**Figure S3.**
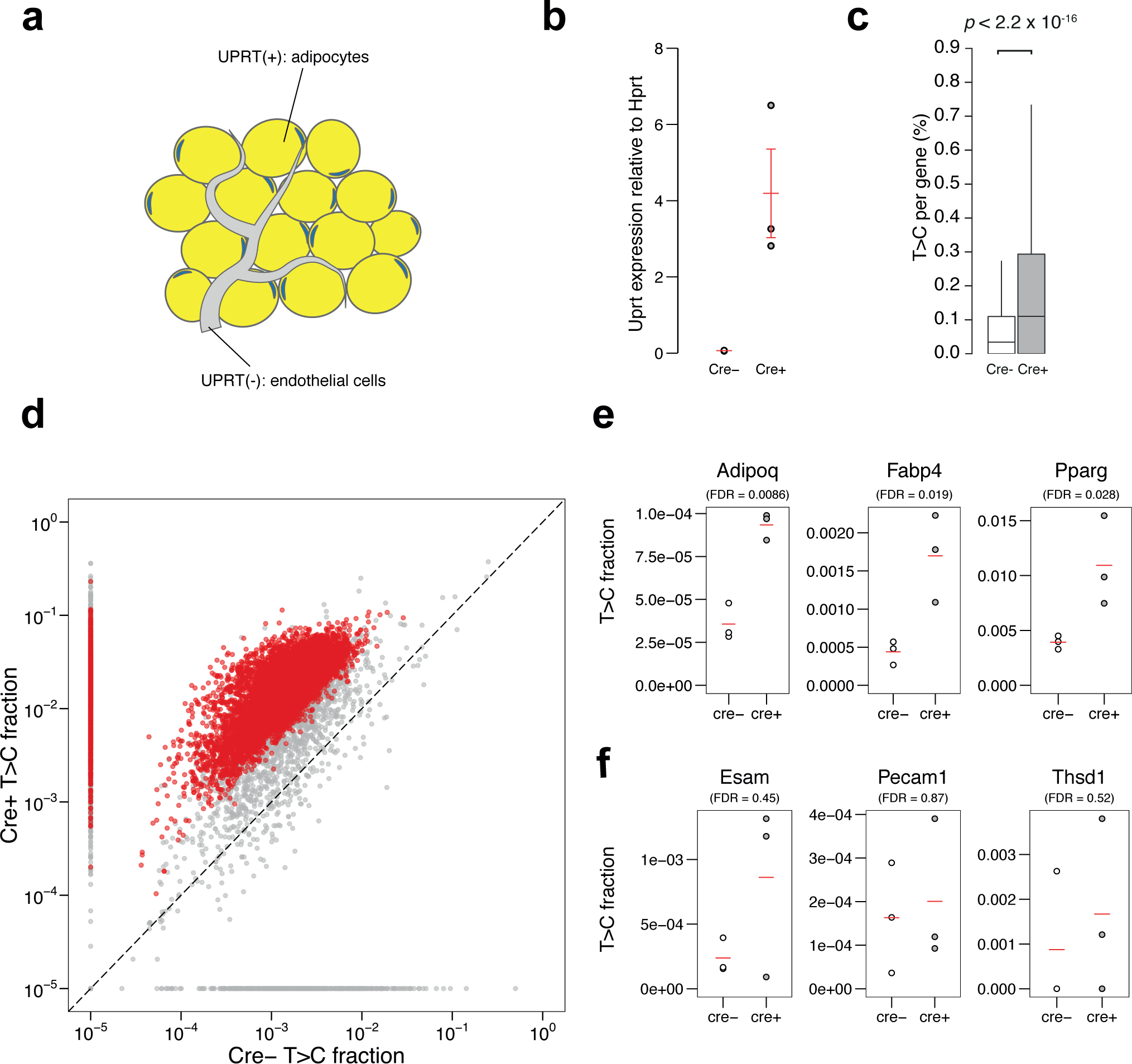
SLAM-ITseq analyses of labelled RNA from the mouse eWAT expressing UPRT in adipocytes. (a) Schematic of UPRT-expressing cells (yellow) and non-UPRT-expressing cells (grey) in the Cre+ mouse intestine. (b) Comparison of UPRT mRNA expression by RT-qPCR in eWAT from the Cre+ and Cre-animals. The red bars indicate the mean expression among biological replicates. (c) Comparison of T>C rate in all T positions sequenced per gene. Data are shown as boxplots. The lower and upper hinges correspond to the first and third quartiles, the middle hinges indicate the median, and the whiskers extend to 1.5 interquartile range from the upper hinges. Outliers are not shown. The p-value obtained by Mann-Whitney *U* test is shown. (d) Average T>C fractions of each gene are plotted. Significantly more labelled transcripts in Cre+ were determined by beta-binomial test and shown as red points (FDR < 0.05). A constant value of 1x10^-5^ was added to the row T>C value when plotting. (e, f) Examples of T>C fraction of known adipocyte-specific genes (e) and endothelial cell-specific genes (f). The red bars indicate the mean T>C fraction of biological replicates (Cre+: n=3, Cre-: n=3).

